# A fine-tuned interplay of three *A. thaliana* UDP glucosyltransferases orchestrates salicylic acid homeostasis

**DOI:** 10.1101/2020.11.02.365221

**Authors:** Sibylle Bauer, Elisabeth Georgii, Birgit Lange, Rafał P. Maksym, Robert Janowski, Birgit Geist, Anton R. Schäffner

**Affiliations:** Institute of Biochemical Plant Pathology, Helmholtz Zentrum München, German, Research Center for Environmental Health, 85764 Neuherberg, Germany; Institute of Structural Biology, Helmholtz Zentrum München, German Research Center for Environmental Health, 85764 Neuherberg, Germany

## Abstract

Salicylic acid (SA) is a central signaling molecule in development and defense, therefore its levels are tightly controlled. One control mechanism is conjugation with sugar moieties by UDP glucosyltransferases (UGTs). In Arabidopsis, UGT76B1, UGT74F1, and UGT74F2 are known to glucosylate SA. We show that these are the main SA UGTs in leaves, since only marginal levels of SA glucosides were found in a triple loss-of-function mutant. Analyzing transcriptomes, metabolite levels, and phenotypes of a full combinatorial set of loss-of-function mutants, we resolved the mutual relationships and the individual roles of these enzymes in SA homeostasis. The strongest gene expression changes were observed for the *ugt76b1 ugt74f1* double mutant, which downregulated developmental genes and most pronouncedly upregulated cell death-related genes. Among the single mutants, *ugt76b1* specifically exhibited increased production of reactive oxygen species, increased resistance to infection, and early senescence. Likewise, higher-order mutations confirmed the dominant role of UGT76B1 in controlling SA levels and thereby the expression of biotic stress response genes. Both UGT74F1 and UGT74F2 affected *UGT76B1* expression. However, while UGT76B1 and UGT74F1 produced SA-2-*O*-β-glucoside, UGT74F2 did not contribute there substantially. Instead, UGT74F2 acted independently of UGT74F1, decreasing steady-state SA levels by producing salicyloyl glucose ester. Remarkably, this did not restrict defense responses. In contrast, UGT74F1 interacted with UGT76B1 in suppressing defense responses. Nevertheless, a benzothiadiazole-triggered defense scenario induced only UGT76B1, whereas UGT74F1 was linked to controlling abiotic stress responses. All three enzymes form a network that, in concert with other UGTs, regulates expression of developmental and stress-related genes.

**One sentence summary:** The salicylic acid glucosylating enzymes of Arabidopsis leaves are crucial for salicylic acid homeostasis and combinatorially impact defense responses and developmental processes.

## INTRODUCTION

Salicylic acid (SA; 2-hydroxybenzoic acid) is an important secondary phenolic compound occurring in a broad range of prokaryotic and eukaryotic organisms. In plants, SA is involved in a multitude of developmental processes and stress responses, playing an essential role during the whole lifespan of the organism (Liu et al., 2015). Both at local and systemic levels, plant resistance to biotrophic pathogens is mediated through SA, and SA biosynthesis is enhanced during plant defense (Song, 2006; Song et al., 2008; Vlot et al., 2009; Zhang et al., 2013; Lu et al., 2016; Dempsey and Klessig, 2017; Vlot et al., 2020). However, high concentrations of endogenous SA have a negative impact on plant growth. Plants with constitutively elevated SA levels exhibit both an enhanced disease resistance and a reduced growth phenotype (Rivas-San Vicente and Plasencia, 2011; Chandran et al., 2014). Therefore, controlling the endogenous free SA levels is crucial for maintaining a tradeoff between growth and defense (Huot et al., 2014).

SA glucosylation suppresses SA signaling and attenuates defense responses (Vlot et al., 2009; Dempsey et al., 2011; Dempsey and Klessig, 2017). Increased SA glucoside levels after pathogen infection might also be storage forms, but reuse has not been shown (Vlot et al., 2009; Vaca et al., 2017). The Arabidopsis glucosyltransferases UGT76B1 (AT3G11340), UGT74F1 (AT2G43840), and UGT74F2 (AT2G43820) can form SA glucosides and thus are possible candidates to influence free SA levels after pathogen infection (Dean et al., 2005; Dean and Delaney, 2008; Song et al., 2008; Noutoshi et al., 2012; George Thompson et al., 2017). There are two types of SA glucosides: Glucosylation at the phenolic group of SA leads to SA-2-*O*-β-D-glucoside (SAG), whereas glucosylation at the carboxyl group leads to salicyloyl glucose ester (SGE), which *in vitro* was only formed by UGT74F2 (Lim et al., 2002; Dean and Delaney, 2008; George Thompson et al., 2017;).

UGT76B1 plays a central role in defense regulation of Arabidopsis. The *ugt76b1* knockout mutations of either *Columbia* (Col) and *Wassilewskija* (Ws) background led to a higher resistance to pathogens (von Saint Paul et al., 2011; Noutoshi et al., 2012). In addition to SA, UGT76B1 glucosylates isoleucic acid (ILA) and N-hydroxypipecolic acid (NHP), which inhibit SA glucosylation and thereby enhance defense in a UGT76B1-dependent manner (Maksym et al., 2018; Bauer et al., 2020a; Bauer et al., 2020b; Holmes et al., 2020; Mohnike et al., 2020). This crucial role of UGT76B1 in balancing the plant defense status raises the question about the roles of UGT74F1 and UGT74F2 in defense or other SA-related processes. Regarding UGT74F1, discrepant resistance phenotypes of the Ws-based *ugt74f1-1* knockout mutant have been reported, ranging from stronger resistance to stronger susceptibility to *Pseudomonas syringae pv tomato* DC3000 (*Pst*) (Noutoshi et al., 2012; Boachon et al., 2014). A defense-regulating effect of UGT74F2 was suggested by a more resistant Col-based *ugt74f2* knockdown line and a UGT74F2 overexpression line with increased susceptibility (Song et al., 2008; Boachon et al., 2014). UGT74F2 glucosylates not only SA but also anthranilate and nicotinate, which is involved in the NAD salvage pathway (Quiel and Bender, 2003; Cartwright et al., 2008; Grubb et al., 2014; Li et al., 2015).

Levels of SA and SA glucosides have previously been studied in single *ugt* mutants. Free SA of *ugt76b1* knockout plants was elevated, but SAG regulation diverged between Col and Ws (von Saint Paul et al., 2011; Noutoshi et al., 2012). Like the resistance phenotype, the reported SA metabolite levels of *ugt74f1-1* knockout plants varied across different studies (Noutoshi et al., 2012; Boachon et al., 2014). Also for the Col *ugt74f2* knockdown mutant contrasting results on SA levels after infection were found (Boachon et al., 2014; Li et al., 2015). Thus, although UGT76B1, UGT74F1, and UGT74F2 can form SA glucosides, their individual roles and interaction in Arabidopsis SA homeostasis remain an open question.

To elucidate functional relationships between the three UGTs in SA glucosylation, this work analyzes a complete set of single, double, and triple loss-of-function mutations of UGT76B1, UGT74F1, and UGT74F2 that were uniformly generated in the Arabidopsis Col background. With the triple mutant containing only marginal levels of SAG and SGE, these three UGTs largely cover the SA glucosylation activity of Arabidopsis Col leaves. We characterize the mutants simultaneously with respect to gene expression profiles obtained by RNA sequencing and with respect to SA metabolite levels measured by liquid chromatography-mass spectrometry (LC-MS). Furthermore, benzothiadiazole (BTH) treatment as well as phenotypic assays were performed to investigate the link to defense reactions.

## RESULTS

### Generation of *ugt* knockout mutations in all combinations

We first generated a full combinatorial set of loss-of-function mutants with the same genetic background, namely Arabidopsis Col. Applying a CRISPR/Cas9-based system for genome editing in Arabidopsis, we generated a *ugt74f1* allele (Clough and Bent, 1998; Fauser et al., 2014). Wild type was transformed with a construct targeting the first exon of *UGT74F1* (Supplemental Methods). Deletion of the A at position 466 of the genomic sequence relative to the ATG translation start resulted in a premature stop codon of the *ugt74f1-2* mutant. Beside the previously studied *ugt74f2-i1a* knockdown line, a *ugt74f2-2* loss-of-function mutant was available (mutant Q153* with a premature stop codon; Quiel and Bender, 2003), which we used after backcrossing and elimination of two additional mutations (*trp1* and *gl1*). Finally, we reused the *ugt76b1-1* loss-of-function allele (von Saint Paul et al., 2011).

Higher-order mutants were generated by genetic crossing if possible. Since *UGT74F1* and *UGT74F2* are positioned in close proximity on chromosome 2, crossing is not feasible. Therefore, the same CRISPR/Cas9 approach as for *ugt74f1* was applied in the *ugt74f2-2* mutant to generate a *ugt74f1-3 ugt74f2-2* double mutant, which contains an insertion of an A after position 466 relative to ATG, leading to another premature stop codon in the first exon of *UGT74F1*. The *ugt74f1-3 ugt74f2-2* double mutant was crossed with *ugt76b1-1* to generate a triple *ugt* mutant. In total, we thus obtained single knockout mutants for UGT74F1, UGT74F2, and UGT76B1 (from now on *ugt74f1, ugt74f2*, and *ugt76b1)* as well as all combinations of knockouts in Col background (Fig. 1A, color key; Supplemental Methods; Supplemental Table 2).

**Figure 1.**
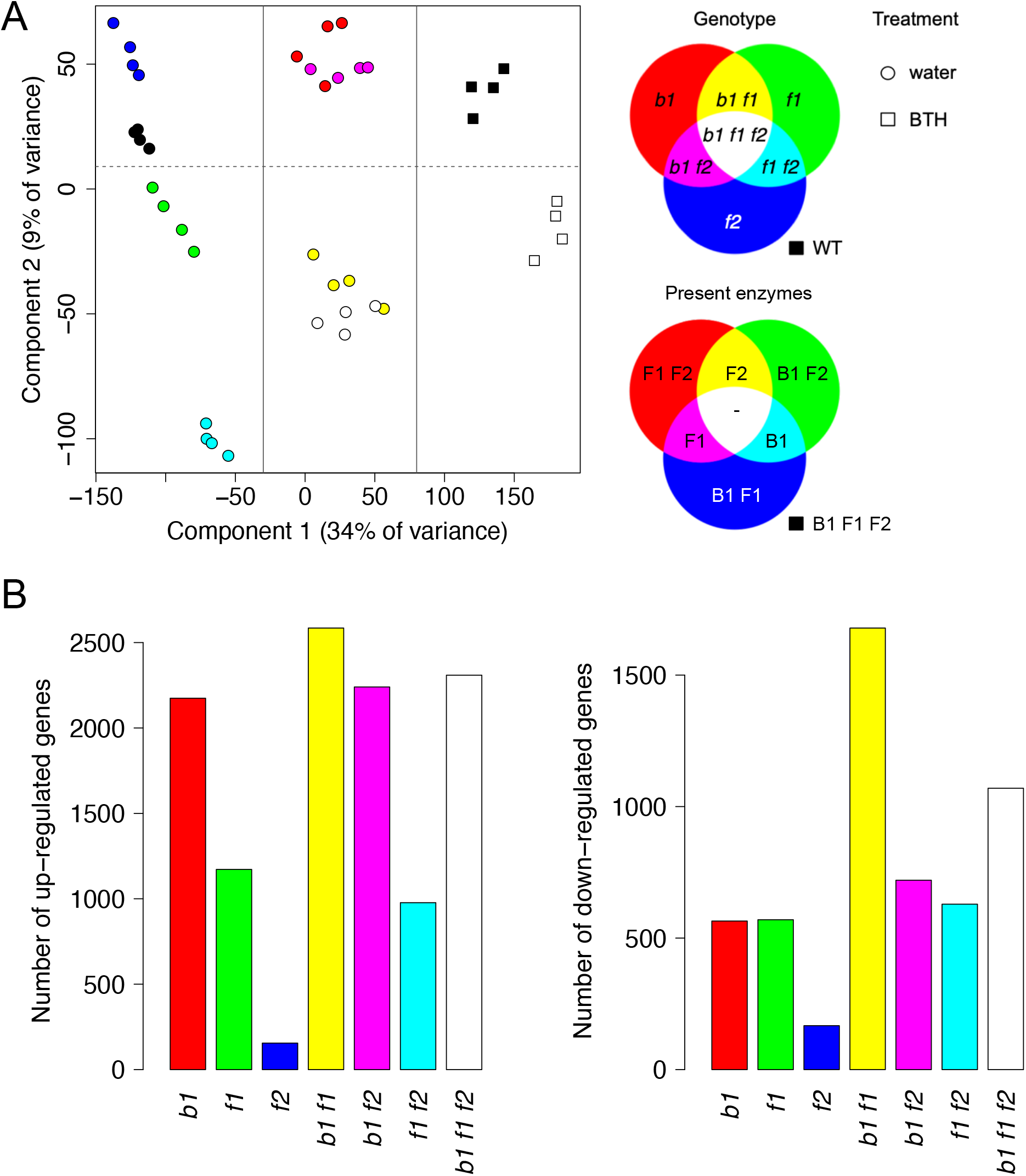
Gene expression profiling of *ugt* mutants. A, Principal component analysis-derived visualization of RNA-seq leaf samples based on gene level read counts normalized as transcripts per million (TPM). Four biological replicates were analyzed for Arabidopsis Col-0 wild-type (WT) plants and a complete set of single, double, and triple knockout mutants for UGT76B1, UGT74F1, and UGT74F2 in Col background. In the color key, the mutated alleles are abbreviated to *b1, f1*, and *f2*, respectively, and the wild-type enzymes to B1, F1, and F2, respectively. WT and *b1 f1 f2* triple knockout plants were additionally treated by the SA analog benzothiadiazole (BTH). The first component separates BTH-treated plants, plants with the *b1* mutation (*b1*, *b1 f1, b1 f2, b1 f1 f2*) and plants without the *b1* mutation (solid gray lines in the plot). The second component separates plants with the *f1* mutation from plants without the *f1* mutation. B, Total number of up- or downregulated genes (adjusted p-value < 0.05 and absolute log_2_ fold change > 1) in mutants compared to WT.

This mutant set enabled a direct comparison of the three UGTs with respect to their biological functions and an assessment of their interplay in SA glucosylation.

### Gene expression profiles of *ugt* knockout mutants

The overall biological impact of *ugt* mutations was investigated at the gene expression level. A leaf RNA-seq analysis yielded gene expression profiles of four biological replicates for each single and combined mutant (Materials and Methods, Supplemental Methods). As a reference, we measured wild-type samples, both under normal conditions and under mimicked stress induced by the treatment with the SA analog BTH. Additionally, the triple knockout mutant was analyzed under BTH stress.

Biological replicates appeared as clusters in the principal component visualization of all gene expression profiles (Fig. 1A). The first component, explaining 34% of the variance, separated BTH-treated plants, plants with the *ugt76b1* mutation, and plants without the *ugt76b1* mutation. The top hundred genes correlating with the first component were significantly enriched in *protein glycosylation* and *transport* functions (p.adj<5.8e-8). The second largest variance component separated plants with the *ugt74f1* mutation from plants without the *ugt74f1* mutation. The top hundred genes correlating with the second component had a significant enrichment for *response to abiotic stimulus* and related functions (p.adj<3.5e-8). Some pairs of mutants belonged to the same cluster in the two-dimensional visualization. That was the case on the one hand for *ugt76b1* and *ugt76b1 ugt74f2*, and on the other hand for *ugt76b1 ugt74f1* and *ugt76b1 ugt74f1 ugt74f2*, confirming the minor impact of the *ugt74f2* mutation.

Differential analysis relative to the wild type revealed that the *ugt76b1 ugt74f1* mutation led to the largest numbers of up- and of downregulated genes among all mutants, whereas the *ugt74f2* mutation had by far the smallest effect (Fig. 1B). Interestingly, the strong effect of the *ugt76b1 ugt74f1* mutation was softened by an additional *ugt74f2* mutation; in particular, the number of downregulated genes was markedly reduced. The genes that were downregulated by *ugt76b1 ugt74f1* and not by *ugt76b1 ugt74f1 ugt74f2* were enriched regarding functions in *extracellular region, tissue development, regulation of growth* and *transcription factor activity* (p.adj<8.9e-8). This indicates that developmental processes were suppressed more strongly in the double mutant than in the triple mutant, suggesting that UGT74F2 inhibits developmental processes and behaves antagonistically to UGT76B1 and UGT74F1.

In summary, the *ugt76b1* and *ugt74f1* mutations had the largest impact on the gene expression profiles and showed distinct characteristics when occurring separately or together. The effect of the *ugt76b1* mutation resembled the effect of BTH treatment.

### SA marker gene expression and defense response of single *ugt* mutants

To further study the biological functions of UGT76B1, UGT74F1, and UGT74F2, we first investigated the single mutants regarding expression changes of SA marker genes (Blanco et al., 2009), which supposedly reflect the levels of non-glucosylated SA. Here, *ugt76b1* had the largest number of upregulated SA marker genes (Fig. 2A). The majority of them were specific to *ugt76b1*, the others were shared between *ugt76b1* and *ugt74f1*, which itself had only very few specific genes. Finally, *ugt74f2* showed upregulation of only four marker genes, all of which were shared with *ugt76b1* and *ugt74f1* but have no immediate role in SA metabolic reactions (AT1G49000, AT3G05660, AT3G48640, AT4G14365).

**Figure 2.**
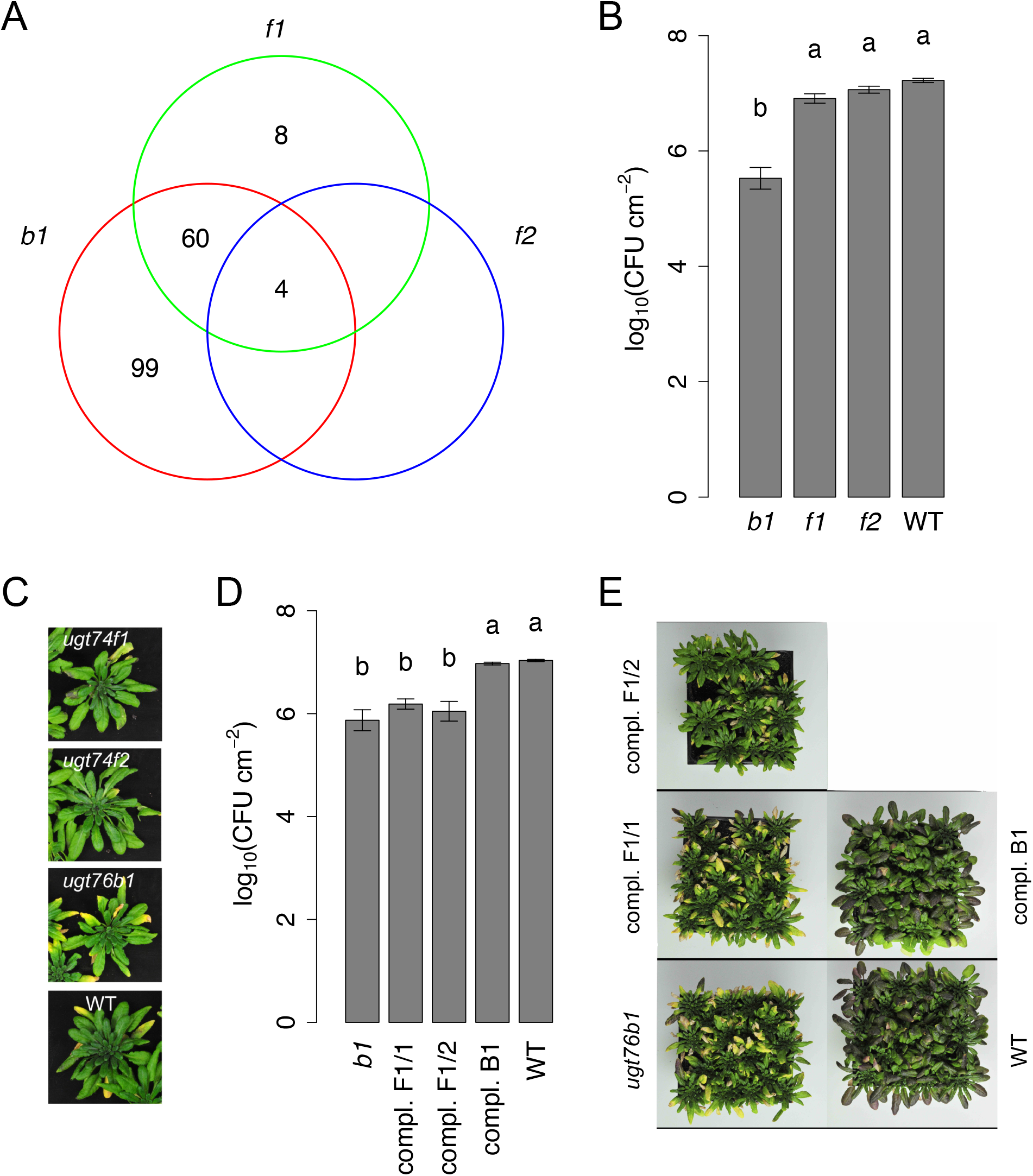
Effects of single *ugt* mutations on SA-related processes. A, Number of SA marker genes other than UGT76B1 that are commonly or specifically upregulated in single mutants relative to the wild type. Abbreviations are the same as in Figure 1. B, Bacterial counts in single mutants and wild type two days after infection with *Pseudomonas syringae pv tomato* DC3000. The graphs shows means ± SE from four biological replicates. Distinct letters indicate significant differences between groups according to one-way ANOVA with Tukey posthoc tests (p<0.05). C, Representative images of eight-week-old plants showing early senescence phenotype of *ugt76b1* compared to *ugt74f1* and *ugt74f2*. D, Bacterial counts of different complementation lines for the *ugt76b1* mutant, using *UGT76B1* 5’ and 3’ regulatory sequences and either the *UGT74F1* coding sequence (compl. F1) or the *UGT76B1* coding sequence (compl. B1), in comparison with *ugt76b1* and wild type. See B for details on the statistical test. E, Eight-week-old rosettes of *ugt76b1*, complementation lines, and wild type.

We then explored whether these differences between single mutants were also visible at the phenotypic level. Indeed, *ugt76b1* plants showed a significantly stronger resistance against *Pst* than *ugt74f1, ugt74f2*, and wild-type plants, which behaved similarly to each other (Fig. 2B). Furthermore, *ugt76b1* exhibited an early senescence phenotype after eight weeks that was not present in *ugt74f1* and *ugt74f2* (Fig. 2C).

Together, the phenotypic results revealed major discrepancies between *ugt76b1* and the other single mutants with respect to defense responses and leaf senescence. SA plays a role in both of these processes (Vlot et al., 2009; Guo et al., 2017). The expression of SA marker genes as well as the overall gene expression profiles substantially differed among all three single mutant lines.

### Spatial expression patterns and structural characteristics of SA UGTs

The observed phenotypic and transcriptomic differences between *ugt76b1* and the other single mutant lines indicate functional differences among the three UGT enzymes. We therefore analyzed whether differences in structure or cellular localization of the enzymes contribute to their specific biological functions.

To complement the whole-leaf transcriptome measurements with information on tissue-specific regulation, we used transgenic lines harboring promoter GFP-GUS reporter fusions. *UGT74F1* and *UGT74F2* were expressed in the leaf vascular tissue. In contrast, *UGT76B1* expression was patchy and spread across the leaf (Supplemental Fig. 1). *UGT76B1* also showed a strong expression in the root tips, which was not detected for *UGT74F1* and *UGT74F2*. This is consistent with the spatial expression patterns reported in the ePlant database (Waese et al., 2017), which additionally reports a strong induction of *UGT74F1* in young leaves. Importantly, the GUS expression patterns did not change when the reporter lines were introgressed into the respective other single mutant backgrounds (Supplemental Fig. 1).

Differences in the active site conformations between UGT74F1 and UGT74F2 are responsible for preferential SAG or SGE formation of the two enzymes, which have an amino acid sequence identity of 77% (George Thompson et al., 2017). A multiple amino acid sequence alignment for UGT74F1, UGT74F2, and UGT76B1 indicated that UGT76B1 had only 27-29% sequence identity to the other two proteins but shared His 18 and Asp 111, which form the catalytic dyad of UGT74F proteins (Supplemental Fig. 2; Gouy et al., 2010; George Thompson et al., 2017). Except for the catalytic dyad and the conserved Asp 366/369, *in silico* structure homology modeling revealed substantial differences in the binding pockets of UGT76B1 and the UGT74F proteins around the SA ligand (Supplemental Fig. 3; Kelley et al., 2015). UGT76B1 has less bulky amino acids and therefore more space at the active site, allowing it to glucosylate other substrates than SA, namely ILA and NHP, which competitively affect UGT76B1’s activity (Maksym et al., 2018; Bauer et al., 2020b; Mohnike et al., 2020).

To evaluate whether these structural and enzymatic differences contribute independently from cellular expression differences to the discrepancy among mutant phenotypes, we constructed a hybrid composed of the *UGT74F1* coding sequence fused with *UGT76B1* 5’ and 3’ regulatory regions and introgressed it in the *ugt76b1* mutant (Supplemental Methods). Two independent hybrid lines generated in this way still showed the same defense and senescence phenotype as the *ugt76b1* mutant, whereas complementation with the complete *UGT76B1* gene sequence restored the wild-type phenotype (Fig. 2D-E).

Thus, the specific structure and enzymatic properties of UGT76B1 play a crucial role in regulating SA-related processes like defense and leaf senescence. This task cannot be performed by the UGT74F1 protein.

### Biological processes altered by single and combined *ugt* mutations

To get insights into functional changes in single and combined *ugt* mutants, a comprehensive Gene Ontology (GO) term enrichment analysis was performed for differentially expressed genes (Supplemental Dataset 1). The GO slim selection of 46 representative biological process terms (Supplemental Methods) was used for an overall visualization, showing that many processes were upregulated across almost all mutants in comparison to the wild type (Fig. 3A). *Cell death* and the (partly overlapping but much more comprehensive) term *response to biotic stimulus* had the largest enrichment for all mutants except for *ugt74f2*, which in fact did not show enrichment with respect to any GO term, underlining the minor impact of UGT74F2 loss on the leaf transcriptome. There was a large overlap of significantly enriched processes between *ugt76b1* and *ugt74f1*, but *ugt76b1* upregulated without any exception many more genes of these processes than *ugt74f1*. The number of upregulated *response to biotic stimulus* genes matches the stronger pathogen resistance of *ugt76b1* (Fig. 2B). *Abscission* was only enriched for *ugt76b1*, including the upregulated gene *SENESCENCE-RELATED GENE 1* (AT1G17020) in consistency with the leaf senescence phenotype observed only for this single mutant (Fig. 2C). *Biosynthetic* and *metabolic process* functions as well as *circadian rhythm* genes were only significantly upregulated by *ugt74f1*, not by *ugt76b1*.

**Figure 3.**
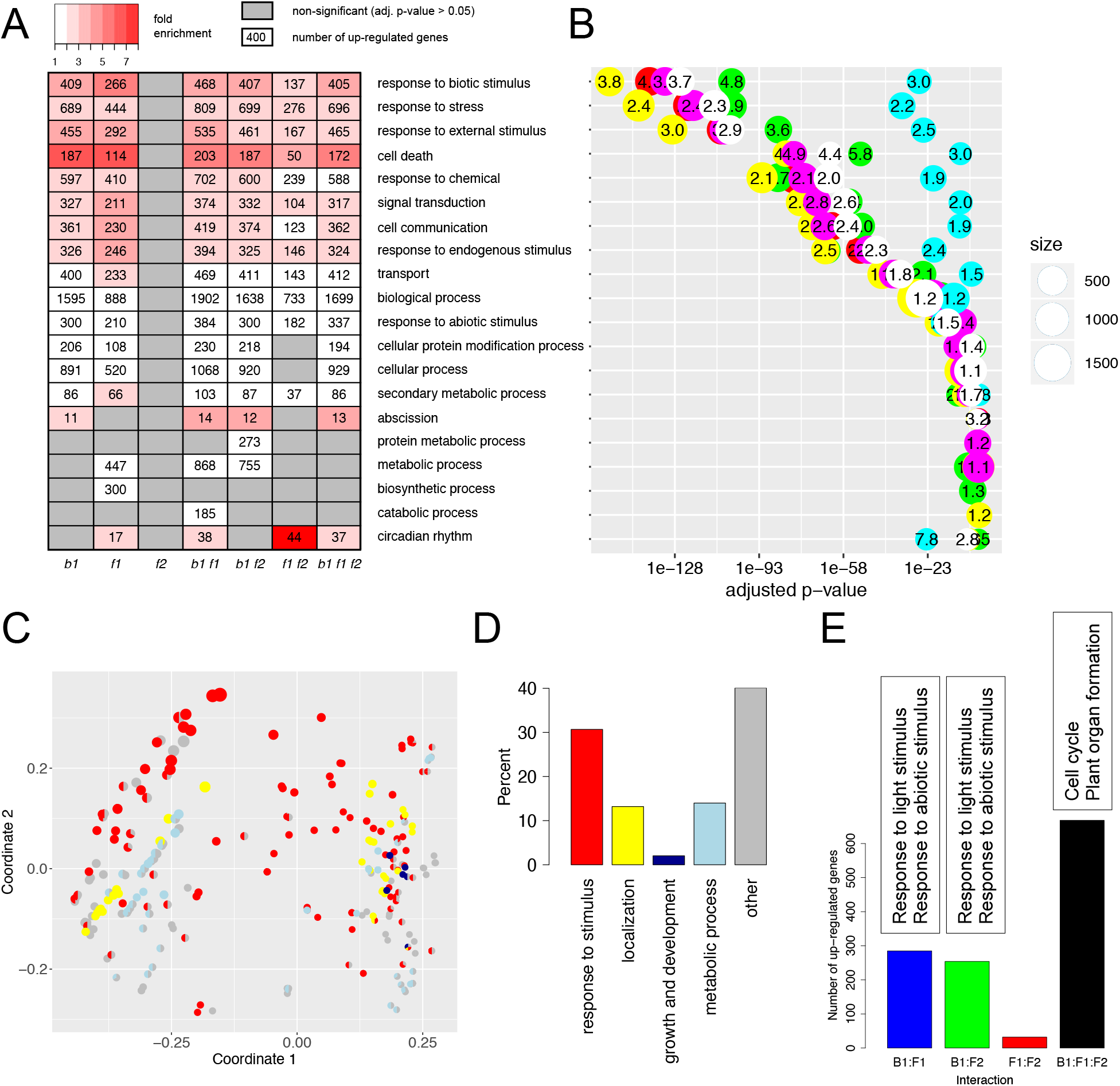
Functional enrichment among genes upregulated by *ugt* mutants. A, Overview of biologically processes upregulated in mutants relative to the WT. The heatmap shows significantly enriched biological process terms that belong to the GO slim selection. B, Adjusted p-value for functional enrichment of the GO terms (A) in each mutant. The mutant colors are taken from the scheme in Fig. 1A. C, Multi-dimensional scaling visualization of all significantly enriched biological process terms of GO (Supplemental Methods). Circle size is proportional to the number of upregulated genes annotated with the term. Circles with larger distance to each other have a smaller number of upregulated genes in common. The color piechart indicates membership of the term in top level categories of different general themes (D). D, Total distribution of themes across all significantly enriched biological process terms. E, Functional interactions among UGT enzymes when functional proteins are present (B1: UGT76B1, F1: UGT74F1, F2: UGT74F2). The bar chart indicates the number of genes showing a significant positive interaction beyond the individual effects of the respective enzymes. The text boxes summarize significantly enriched functions for these gene groups.

Among the *ugt* mutants, *ugt76b1 ugt74f1* had the largest number of upregulations in all enriched processes and went beyond the effects of the *ugt76b1* and *ugt74f1* single mutants. This is also reflected in the strong significance of the adjusted enrichment p-values for *ugt76b1 ugt74f1*, outperforming the other mutants (Fig. 3B). The *ugt76b1 ugt74f1* mutant was enriched in all the processes that had shown up for either the *ugt76b1* or the *ugt74f1* mutant, with one interesting exception: Instead of *biosynthetic processes*, *catabolic processes* were activated. In fact, biosynthesis was downregulated along with growth and development (see below). The *ugt74f1 ugt74f2* double mutant upregulated only approximately half of the *response to biotic stimulus* or *cell death* genes that were upregulated by the *ugt74f1* single mutant, supporting a counteracting effect of the *ugt74f2* mutation. Similarly, the *ugt76b1 ugt74f1 ugt74f2* triple mutant had less upregulated genes in these processes than the *ugt76b1 ugt74f1* double mutant.

In total, four hundred GO terms were enriched among upregulated genes of the triple mutant (Supplemental Dataset 1). We visualized the biological process terms according to their overlap of upregulated genes and grouped them according to their broad theme (Fig. 3C). Clearly, the majority of terms and also the terms with the largest numbers of upregulated genes were related to stress responses (Fig. 3D). The top most significantly enriched terms were *defense response* and *systemic acquired resistance* (SAR). Furthermore, *SA biosynthetic process, response to SA*, and *regulation of cell death* were at the top of the list (p.adj<2.0e-65), followed by several different *localization and signaling* processes, and *regulation of reactive oxygen species* (ROS) *metabolic process* (p.adj<3.0e-47). The term ranking was highly similar to *ugt76b1*, showing the dominance of this mutation.

*Responses to abiotic stimulus, temperature*, and *water deprivation* also showed an enrichment among upregulated genes of the triple mutant. In contrast, *response to light stimulus* was enriched among downregulated genes, making up the main fraction of its 149 downregulated genes in *response to abiotic stimulus*. Indeed, upregulation of *response to light stimulus* depended on the presence of UGT76B1 and UGT74F1 or UGT74F2 (Fig. 3E and next subsection). The *ugt76b1 ugt74f1* double mutant downregulated even more light-responsive genes and – in contrast to the triple mutant – many processes related to development and growth (Fig. 4A). The *ugt74f1* single mutant downregulated some developmental processes and secondary metabolism, but not growth.

**Figure 4.**
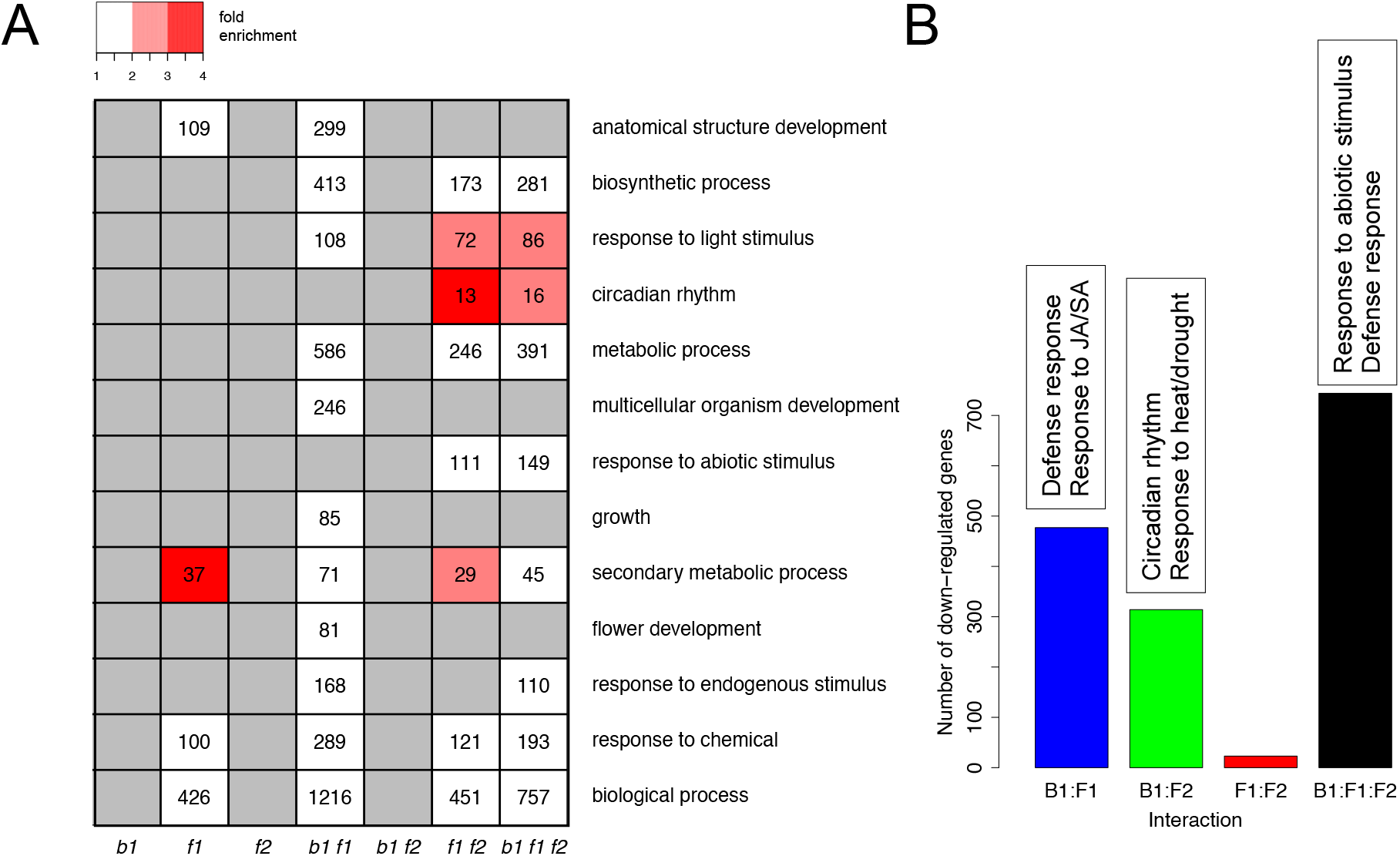
Functional enrichment among genes downregulated by *ugt* mutants. A, Heatmap of significantly enriched GO slim terms. B, Negative functional interactions among UGT enzymes. The bar chart indicates the number of genes showing a significant negative interaction of the respective enzymes. The text boxes summarize significantly enriched functions for these gene groups.

In summary, *ugt76b1 ugt74f1* was the most affected mutant in our combinatorial set, showing tremendous expression upregulation in biotic stress response and cell death genes and severe downregulation of development and growth processes compared with the wild type.

### Functional interactions of UGT enzymes

The comprehensive set of mutants for the three enzymes UGT76B1, UGT74F1, and UGT74F2 allowed us to study their functional interaction effects on the expression of other genes. The *ugt76b1 ugt74f1 ugt74f2* triple mutant forms the reference based on which the effect of adding one or several functional proteins will be evaluated.

The presence of UGT76B1 alone (i.e. *ugt74f1 ugt74f2*) led to an upregulation of *transcription factor activity, response to JA, JA metabolism*, and *response to water deprivation* or *ABA*, and to a downregulation of *defense response, SAR* and *SA biosynthesis*. UGT74F1 induced an upregulation of light-related responses and a downregulation of *circadian rhythm* and temperature and drought responses. Co-occurrence of both enzymes had positive interaction effects on light-related and general abiotic responses as well as *glucosyltransferase activity* (Supplemental Dataset 2, Fig. 3E), whereas it negatively affected *defense* and responses to both JA and SA (Fig. 4B). Upregulation effects were dominated by UGT74F1, downregulation effects were dominated by UGT76B1.

UGT74F2 presence was characterized by a repression of several transport-related processes (Supplemental Dataset 2). Interestingly, UGT74F1 and UGT74F2 showed almost no interaction (Figs. 3E and 4B), suggesting independent modes of action. In contrast, when UGT74F2 co-occurred with UGT76B1, approx. 250 genes displayed a positive interaction effect and approx. 300 genes displayed a negative interaction effect. The former gene set was enriched in mostly light-related abiotic stress responses, the latter in *circadian rhythm* and heat- and drought-related abiotic stress responses. Positive interaction effects were very similar when either UGT74F2 or UGT74F1 joined UGT76B1. With the interaction of all three enzymes growth and development processes were promoted and responses to both biotic and abiotic stresses were downregulated (Figs. 3E and 4B).

In summary, UGT76B1 showed strong interaction effects with both other enzymes.

### SA metabolite levels of *ugt* mutants

To further elucidate the functional interactions between the UGT enzymes, we were interested how their combinatorial presence or absence influences global levels of their common substrate SA and their glucosylation products SAG and SGE. For that purpose, metabolite levels were determined by LC-MS. Under control conditions, the *ugt74f1, ugt74f2*, and *ugt74f1 ugt74f2* mutants showed SA levels similar to the wild type, whereas all mutants containing the *ugt76b1* mutation showed significantly enhanced SA levels (Fig. 5A), suggesting that UGT76B1 controls SA levels. A significant increase of SAG levels relative to the wild type was only observed for *ugt76b1 ugt74f2*. SGE was significantly enhanced in *ugt76b1* and *ugt76b1 ugt74f1*. Clearly, enhanced production of SAG and SGE occurred only in the absence of UGT76B1 and depended on UGT74F1 and UGT74F2, respectively.

**Figure 5.**
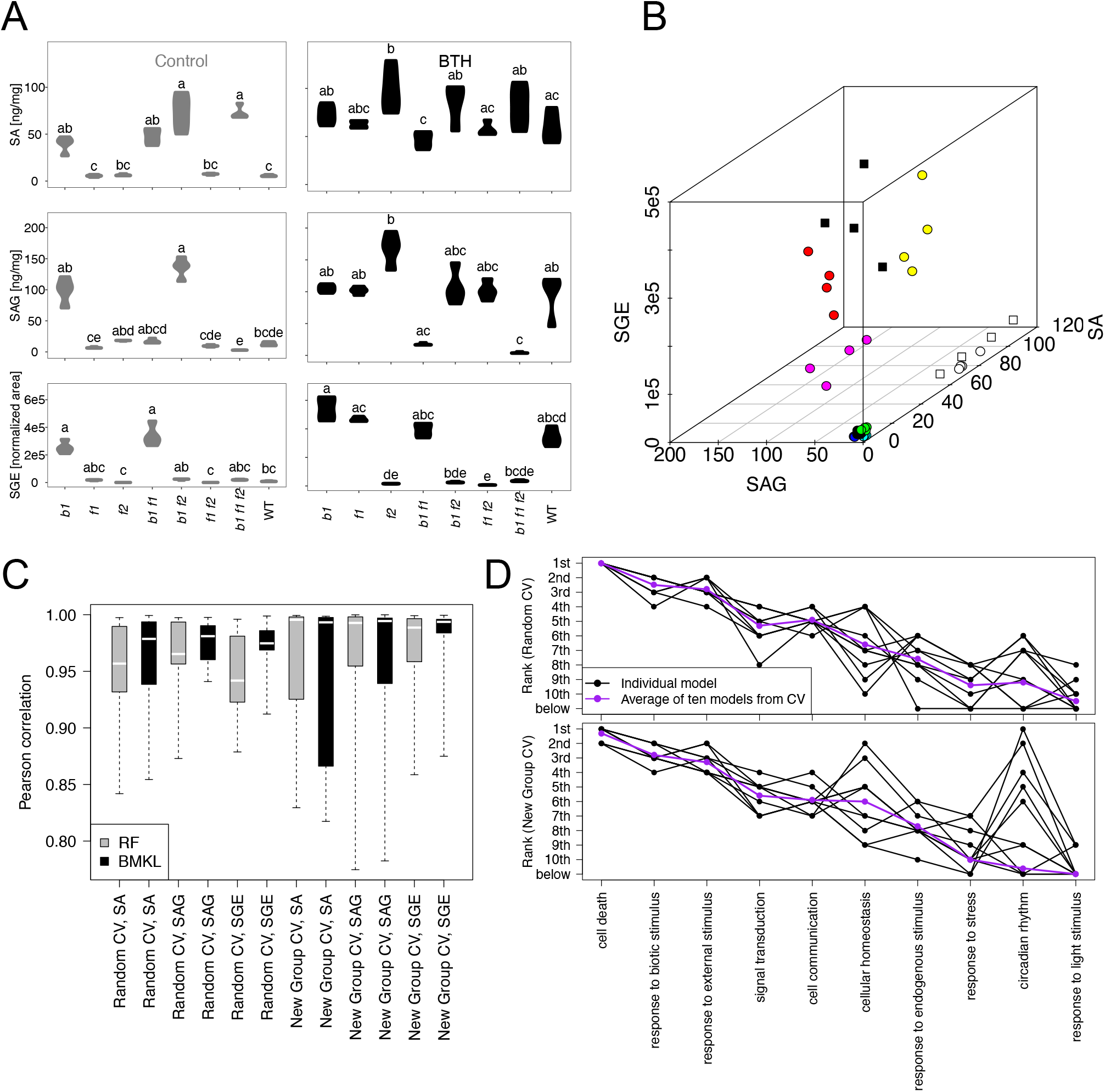
Changes in SA metabolite levels and associated gene expression changes of *ugt* mutants. A, Violin plots for levels of three SA metabolites measured by LC-MS for *ugt* mutants and BTH treatment. Groups marked by distinct letters are significantly different (Kruskal-Wallis test with Dunn’s posthoc test p<0.05). B, SA metabolite levels of the biological samples that were used for RNA-seq analysis. Color scheme is identical to that of Fig. 1A. C, Performance of predicting SA metabolite levels from gene expression data. Two modeling approaches were performed, Random Forests (RF) and Bayesian Multiple Kernel Learning (BMKL). Both approaches were applied on the same training and test data sets. For the training data, gene expression variables were standardized and SA metabolite data were centered. The respective mean and standard deviation values derived from the training data were used to normalize the test data. Two types of tenfold cross-validation (CV) were performed, where the test set consisted either of a previously unseen random subset of samples (Random) or an entirely new biological group (New Group). The y axis indicates the correlation between true and predicted differences of the test samples to the training samples. Boxes show 25% and 75% quantiles, the white line represents the median and the whiskers indicate the extreme values across the ten folds. D, Importance ranking of GO slim terms obtained by BMKL. Both CV runs yielded the same top ten GO slim terms, here sorted by average rank of the New Group CV.

After treatment with BTH, wild-type levels of all three compounds went up compared with control conditions (Fig. 5A). Remarkably, production of SAG was reduced to marginal levels in the triple mutant, significantly below the wild type, suggesting that no other enzymes take over this function. SGE levels of *ugt74f1 ugt74f2* dropped significantly below the wild-type concentration and became undetectable, showing that UGT76B1 cannot produce it. SA levels of *ugt74f2* were significantly above those of the wild type and *ugt76b1 ugt74f1*, indicating that UGT74F2 restricts the free SA level under BTH treatment.

Taken together, while all three SA-related compounds are induced by BTH, each of them is regulated in a distinct manner by a combination of functional UGTs. UGT74F1 and UGTF2 promote the production of SAG and SGE, respectively, whereas UGT76B1 has a main role in keeping free SA levels down.

### Gene expression changes associated with SA homeostasis

Given the complex response of SA-related compounds to *ugt* mutations and BTH treatment, we were interested whether their levels could be predicted from gene expression profiles, thus revealing the most relevant associated biological processes. For that purpose we applied two methods that performed best in prediction of several quantitative response variables (Costello et al., 2014): Bayesian Multiple Kernel Learning (BMKL) and Random Forests (RF). The measurements of SA, SAG, and SGE from the samples used for the RNA-seq analysis showed the same trends as discussed in the previous section for the full LC-MS dataset, and biological replicates co-occurred in clusters (Fig. 5B).

The samples of the dataset were divided into ten groups to perform cross-validation, each time keeping one group as a test set and taking the remaining samples for training. In addition to the standard cross-validation based on random splitting, we considered the more challenging task of testing predictions for each biological group, i.e. genotype-treatment combination, when training was performed only with the other biological groups. For a fair performance evaluation in spite of the changing output value range of training data across the cross-validation folds, we took the correlation between true and predicted differences of the test samples to the training samples as a quality assessment criterion.

Both methods performed well, achieving a correlation of more than 0.8 in almost all cases and median values greater than 0.94 across the cross-validation folds for all SA-related compounds and prediction tasks (Fig. 5C). This indicates that reliable relationships between gene expression and compound levels were detected. For the more difficult task of predicting a new biological group, the performance showed larger variance and range of extreme values than for predicting unseen samples of known biological groups. Overall, the predictions of BMKL tended to be more robust, with greater or comparable median values relative to RF. Therefore, we analyzed the BMKL models in more detail to get insights into potential biological relationships. For each cross-validation fold, BMKL learned a single model for all three compounds. In contrast to RF, BMKL does not yield importance weights for single genes but for predefined groups of genes, limiting the number of parameters to learn. We used the 46 GO slim biological process terms as gene groups. Remarkably, the top ten highest-ranking GO slim groups were identical for both cross-validation tasks (Fig. 5D). Both times the most predictive group was *cell death* (GO:0008219). It received the top weight in each random cross-validation fold and seven times in the group cross-validation.

Among the *cell death* genes, the top candidates at the individual gene level were investigated for both methods. The BMKL analysis was repeated defining each *cell death* gene as its own group. This yielded as top genes *MPK3* (*MITOGEN-ACTIVATED PROTEIN KINASE 3*; AT3G45640) and *SOT12* (*SULFOTRANSFERASE 12*; AT2G03760). *MPK3* was significantly upregulated by *ugt76b1, ugt76b1 ugt74f1, ugt76b1 ugt74f2*, and BTH-treated wild type compared with wild-type control, matching the regulation pattern of SGE and SAG. *SOT12* is known to respond to SA and to form SA sulfonate. For RF, the top-ranked *cell death* genes were *RIN4* (*RPM1 INTERACTING PROTEIN 4*; AT3G25070), associated to bacterial defense, and the ozone-responsive ARM repeat superfamily protein AT3G02840 (Berardini et al., 2015). *MPK3* also appeared among the top 15 genes. However, for both methods the prediction capacity dropped when focusing on *cell death* genes, demonstrating the benefit of wider gene profiles for robust predictions.

In summary, it was possible to relate UGT-dependent changes in levels of SA-related compounds to gene expression profiles, and the most predictive gene groups were *cell death* and, partly overlapping but much broader, *response to biotic stimulus*.

### ROS formation related to SA homeostasis

The prediction analysis revealed *cell death* as the top gene group whose expression levels were associated with the levels of SA, SAG, and SGE in Arabidopsis leaves. In addition to the common prediction factors for all three compounds identified in the previous section, we extracted for each individual compound the single differentially expressed gene from the *cell death* category that correlates best with the compound level changes across samples from all UGT genotype combinations and treatments (Fig. 6A).

**Figure 6.**
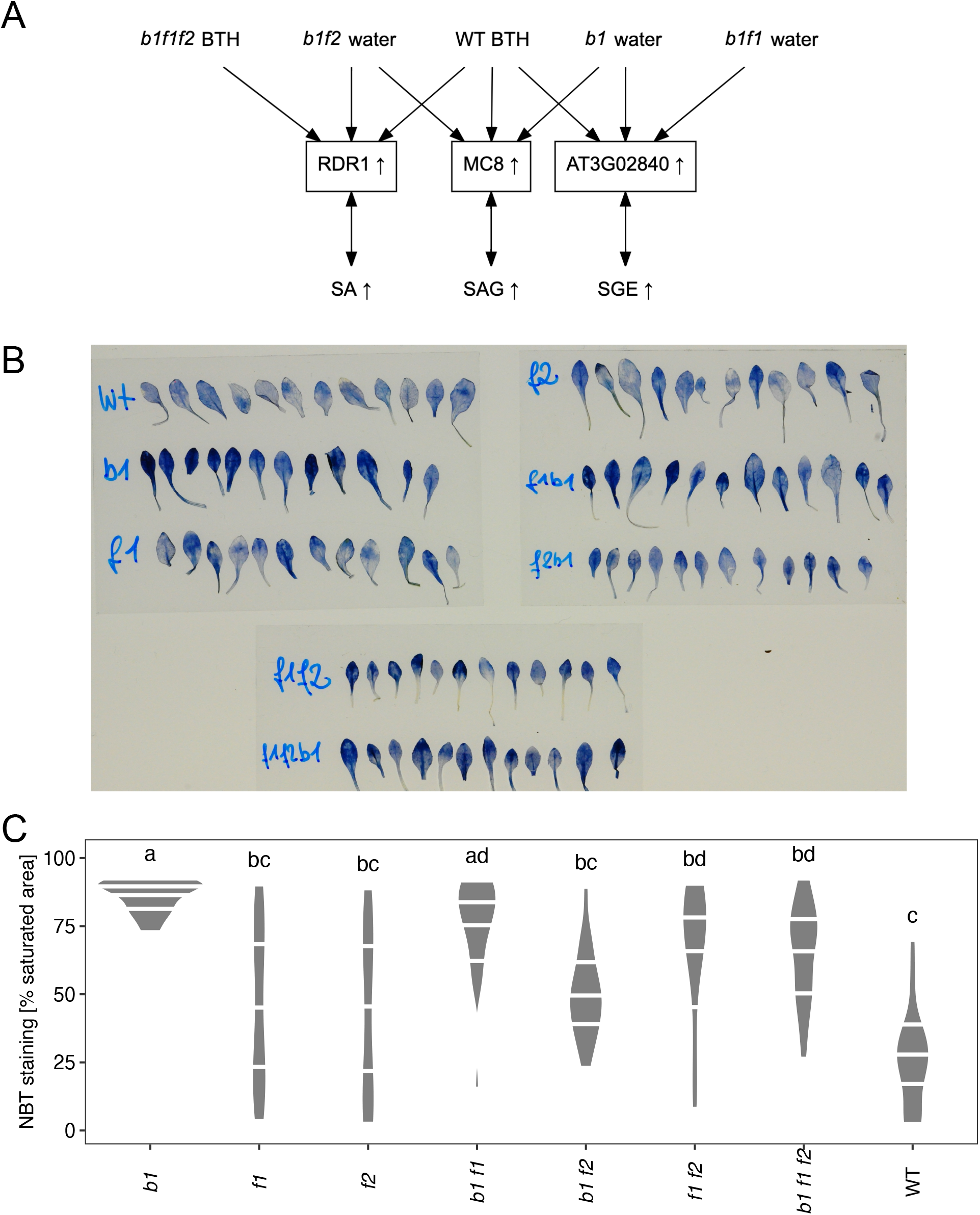
Relationship of SA metabolism to ROS. A, Top cell death genes positively associated with levels of each SA metabolite, obtained from correlation analysis with expression data of all differentially expressed genes in the top category from Fig. 5D. Genes are connected to the three conditions (composed of genotype and treatment) where they showed the largest significant fold change compared to WT water. These are consistent with significant increases observed in metabolite levels (Fig. 5A). B, Images of wild-type and *ugt* mutant leaves after infiltrating two-week-old *A. thaliana* plantlets from liquid culture with nitroblue tetrazolium (NBT). C, Violin plot of NBT staining relative to leaf area, with white horizontal lines for 25, 50, and 75 % quantiles. Groups marked by distinct letters are significantly different (Kruskal-Wallis test with Dunn’s posthoc test p<0.05).

For SA, we obtained *RDR1* (*RNA-DEPENDENT RNA POLYMERASE 1*; AT1G14790) as the top associated gene. *RDR1* expression matched well the pattern of measured SA levels: it was upregulated in all four mutants with the *ugt76b1* mutation and for all BTH-treated groups compared with the wild-type control and had the largest fold changes for BTH-treated groups and *ugt76b1 ugt74f2* (Fig. 5A-B). RDR1 is involved in response to virus infection and induced by SA (Campos et al., 2014; Cao et al., 2014). The top gene associated with SAG was *MC8* (*METACASPASE 8*; AT1G16420), which has been linked to programmed cell death induced by ROS (He et al., 2008). In consistency with the SAG measurements, *ugt76b1 ugt74f2, ugt76b1*, and BTH-treated wild type showed the strongest significant upregulations. The top gene for SGE was AT3G02840, sharing the strong upregulation in *ugt76b1 ugt74f1*, BTH-treated wild type, and *ugt76b1*. This gene is most strongly expressed in senescent leaves (Berardini et al., 2015; Klepikova et al., 2016) and is also induced by ROS (Inzé et al., 2012).

Both *MC8* and AT3G02840 point towards a close relationship between SA-related compounds and ROS, suggesting that ROS levels vary among the *ugt* mutants. In general, ROS signaling, including mitogen-activated protein kinases such as MPK3, plays an important role in cell death (Van Breusegem and Dat, 2006), which we identified as the most relevant gene functional category for SA metabolites. To investigate ROS levels in our mutant collection, we directly tested O_2_^-^ radical formation of the *ugt* mutants by nitroblue tetrazolium (NBT) staining (Fig. 6B-C). Here, *ugt76b1, ugt76b1 ugt74f1, ugt74f1 ugt74f2*, and the triple mutant showed enhanced signals relative to the wild type.

Thus, we conclude that ROS formation is affected by all three UGT enzymes but is not directly associated with levels of SA, SAG, or SGE.

### Relationship of UGT76B1, UGT74F1, and UGT74F2 to other UGTs

Finally, we investigated more closely the relationship of UGT76B1, UGT74F1, and UFT74F2 to the other UGTs (Paquette et al., 2003), to be aware of any compensatory or co-regulatory effects among related enzymes. Many UGTs were not only differentially expressed in at least one mutant or treatment condition but also individually separated specific mutant or treatment groups from all the other groups via a specific expression threshold (Fig. 7A, Supplemental Methods). Expression of these genes was either positively or negatively associated with specific mutation or treatment conditions. Together, they form a network, revealing also relationships between conditions (Fig. 7A).

**Figure 7.**
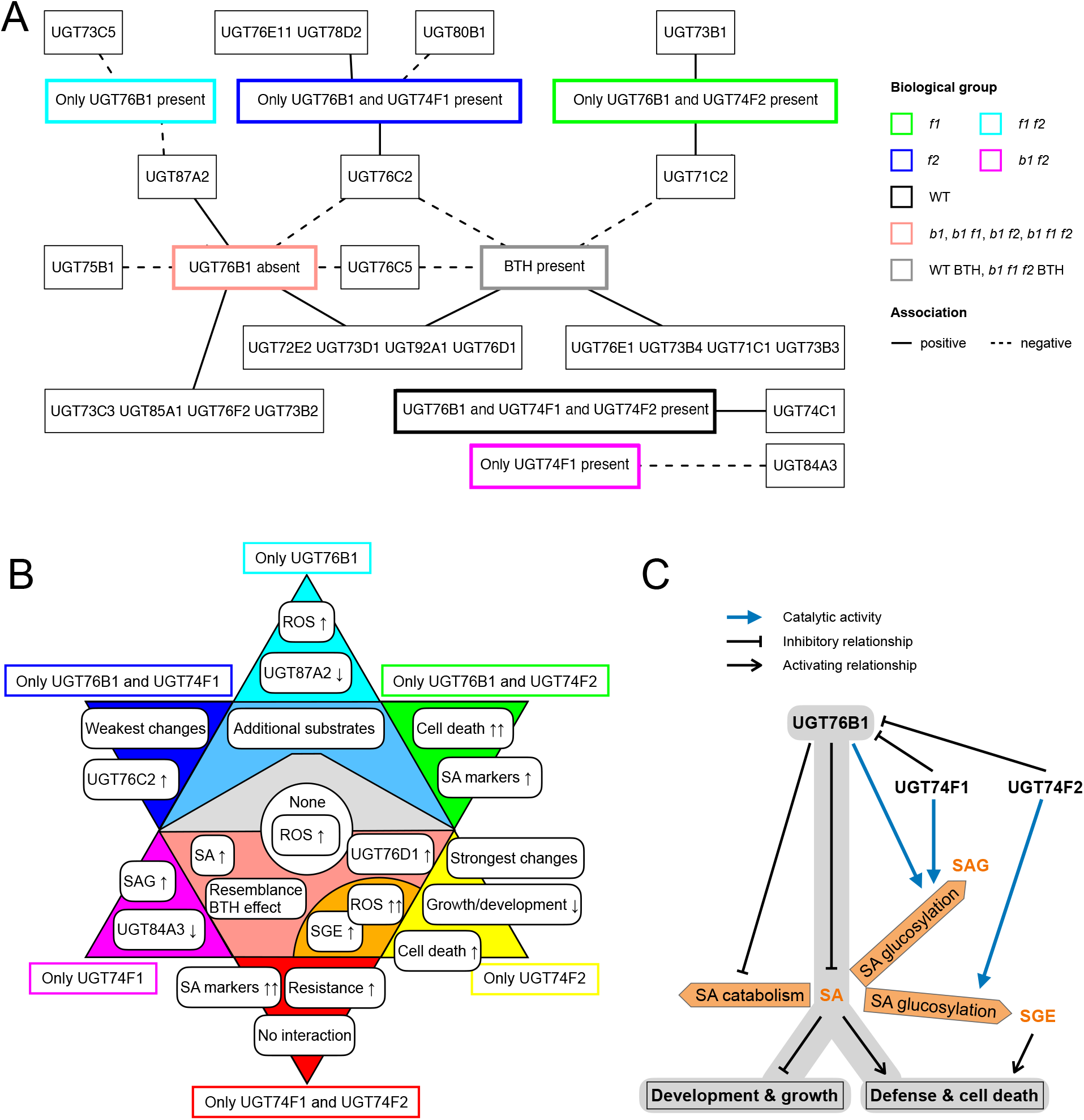
Biological functions and relationships of UGT enzymes. A, Effects of presence or absence of UGT76B1, UGT74F1, and UGT74F2 as well as BTH on the expression of other *UGT* genes. The graph shows differentially expressed genes that completely separate the given groups from the other samples (Methods). Positive associations are marked as solid lines and indicate that expression exceeds a threshold, negative associations are marked as dashed lines and indicate that expression falls below a threshold under the given conditions. B, Summary of main findings for different presence/absence constellations of UGT76B1, UGT74F1, and UGT74F2. Individual mutants are marked in the same colors as in Fig. 1A, with the triple mutant at the center and double and single mutants in the periphery. Groups of mutants are represented by colored shapes that include all mutants whose borders are touched. Round rectangles represent major characteristics of each group. A downward arrow represents a downregulation and an upward arrow an upregulation; a double arrow indicates the strongest effect among all noted cases. For instance, the largest fold enrichment value for *cell death* genes was found in the *f1* mutant (*Only UGT76B1 and UGT74F2*). C, Model of activating and inhibitory relationships among UGT enzymes, SA metabolites, and biological processes. The main metabolic reactions discussed here are depicted in orange: SA synthesis and formation of the glucosides SAG and SGE.

The most obvious finding is that the effect of BTH presence resembles the effect of UGT76B1 loss, irrespective of further losses. Already observed from the PCA of whole gene expression profiles (Fig. 1A), the relationship is here established solely via UGT expression. BTH presence and UGT76B1 loss shared for instance a distinct increase of *UGT76D1* and a distinct decrease of *UGT76C2* and *UGT76C5*. UGT76D1 is known to glucosylate catabolic products of SA (Huang et al., 2018). *UGT85A1* expression was also tightly co-regulated with UGT76B1 loss and the corresponding increases in SA levels (Supplemental Dataset 3, Fig. 5A). Upregulation of *UGT87A2* indicated the loss of UGT76B1, whereas its downregulation was related to the sole presence of UGT76B1 (i.e. loss of UGT74F1 and UGT74F2). Similarly, *UGT71C2* revealed BTH treatment by lower expression and loss of only UGT74F1 by enhanced expression. Some *UGT*s were specifically associated with one mutant, for instance UGT84A3 expression was downregulated by the *ugt76b1 ugt74f2* mutant, where only UGT74F1 was present.

In total, the gene expression profiles imply a sophisticated, fine-tuned interplay of *UGT* genes. Specific groups of *UGT* genes were transcriptionally activated or deactivated in response to combinatorial mutations or stress treatment.

## DISCUSSION

The comprehensive set of loss-of-function mutants of Arabidopsis SA glucosyltransferases revealed interesting combinatorial effects of the presence or absence of UGT76B1, UGT74F1, and UGT74F2 (Fig. 7B). First of all, our mutant analyses consistently confirm the pivotal function of these enzymes in the formation of SA glucosides. Since the triple mutant had the lowest level of SAG among all genotypes, both under control conditions and BTH treatment (Fig. 5A), all three UGT enzymes contribute to SAG formation *in vivo*. Furthermore, our data support the major role of UGT74F2 in SGE formation (Lim et al., 2002; Noutoshi et al., 2012; George Thompson et al., 2017): All mutants that lost UGT74F2 did not show the BTH-induced increase of SGE that was observed for the wild type and the remaining *ugt* mutants (Fig. 5A). Thus, significant SGE formation was only possible when UGT74F2 was present (Fig. 7B).

Beyond that, specific functions of UGT76B1 and UGT74F1 in SA homeostasis were revealed. UGT76B1 was essential to restrict the levels of free SA under normal conditions; whenever UGT76B1 was missing, SA levels increased (Figs. 5A and 7B). The metabolite data suggest that this control does not happen via SAG formation. Under normal conditions, SAG levels could only rise substantially when UGT74F1 was present and UGT76B1 was absent (Figs. 5A and 7B), whereas both UGT74F1 and UGT76B1 contributed to increased SAG production under BTH treatment (Fig. 5A). Indeed, the structure analysis (Supplemental Fig. 3) and previous studies (Bauer et al., 2020a; Bauer et al., 2020b; Mohnike et al., 2020) propose a unique role of UGT76B1 in integrating additional immune-modulatory molecules that influence SA homeostasis. Thereby, UGT76B1 regulates the plant defense status. Accordingly, the expression of the SA synthesis gene *ICS1* (*ISOCHORISMATE SYNTHASE 1*; AT1G74710) was upregulated by all mutants lacking UGT76B1, independently of the presence or absence of the other two SA glucosyltrasferases (Supplemental Table 3, Supplemental Dataset 3). Neither UGT74F1 nor UGT74F2 could replace UGT76B1 in the function of controlling SA levels (Fig. 5A), even if UGT74F1 was expressed under the control of the regulatory sequences of UGT76B1 (Fig. 2D-E). Thus, the unique properties of the UGT76B1 enzyme rather than its expression pattern are responsible for its distinct function. Accordingly, *UGT74F1* was not upregulated at the loss of UGT76B1.

Remarkably, *UGT76B1* was immensely upregulated upon BTH treatment of the wild type, whereas *UGT74F1* remained unchanged (Supplemental Dataset 3). This is in agreement with the role of UGT76B1 as the primary SA signaling attenuator during pathogen stress (Bauer et al., 2020b; Holmes et al., 2020; Mohnike et al., 2020). In contrast, combined drought-heat stress led to an upregulation of *UGT74F1* and no change in the expression of *UGT76B1* (Georgii et al., 2017). Since SA signaling also plays a role in mediating abiotic stress responses (Rivas-San Vicente and Plasencia, 2011), UGT74F1 may be the specific attenuator under these conditions. Consistently, loss of UGT74F1 led to an enriched upregulation of drought- and heat-responsive genes (Supplemental Dataset 1), and addition of UGT74F1 to the triple mutant downregulated these as the top enriched groups (Supplemental Dataset 2).

Apart from these unique functions, UGT76B1 and UGT74F1 share several commonalities. Although *ugt74f1* showed a smaller number of induced SA marker genes than *ugt76b1* (73 vs. 172), 89% of the genes were the same as for *ugt76b1* (Fig. 2A), suggesting that UGT76B1 and UGT74F1 have several common downstream processes. These mainly include the control of biotic stress responses and cell death, with *cell death* showing the highest fold enrichment after loss of UGT74F1 (Fig. 3A). Moreover, the expression data even indicate a functional interaction of UGT76B1 and UGT74F1 in the control of defense response including responses to JA and SA (Fig. 4B) but also in the control of abiotic stress responses (Supplemental Dataset 2), putatively to promote growth and development processes under non-stress conditions. Consistently, gene expression related to growth, light response, biosynthesis, and plant development significantly decreased in the double mutant where both UGT76B1 and UGT74F1 were lost, whereas expression related to response to stress, response to biotic stimulus, and cell death increased, with the largest number of affected genes among all the mutants (Figs. 3A, 4A, and 7B).

The largest number of transcriptionally upregulated transport genes was also found in *ugt76b1 ugt74f1*, reflecting the importance of transport in defense processes, including the transport of defense signals to promote systemic defense reactions (Park et al., 2007; Waszczak et al., 2018; Maruri-López et al., 2019). Systemic acquired resistance was among the top ten enriched GO terms for the genes that were upregulated in *ugt76b1 ugt74f1* (Supplemental Dataset 1). *UGT74F1* was most strongly expressed in the leaf vascular tissue (Supplemental Fig. 1). *UGT76B1* was also expressed in the leaf vascular tissue and overall upregulated after the loss of UGT74F1 (Supplemental Table 1). Therefore, the leaf vascular tissue as transportation hub could be important for the suppression of defense responses by the interaction of UGT76B1 and UGT74F1 under non-stress conditions (Fig. 4B). When SA was enhanced under stress treatment, both enzymes contributed to converting SA to SAG and largely compensated for the loss of each other (Fig. 5A). However, UGT74F2 did not take over this function to a significant degree.

Instead, UGT74F2 was clearly responsible for SGE formation (Figs. 5A-B and 7B). Thus, UGT74F1 and UGT74F2 focus on different reactions that do not interfere with each other. Accordingly, gene expression profiles revealed almost no functional interaction between UGT74F1 and UGT74F2 (Figs. 3E, 4B, and 7B). However, both UGT74F1 and UGT74F2 interacted with UGT76B1. In fact, UGT76B1 levels significantly increased more than twofold when UGT74F1 or UGT74F2 was lost (Supplemental Table 1, Supplemental Dataset 3), whereas this was not the case the other way round. This suggests that UGT74F1 and UGT74F2 independently restrict *UGT76B1* expression via their enzymatic activity (Fig. 7C). In contrast, processes related to the other substrates of UGT74F2, nicotinate and anthranilate (Quiel and Bender, 2003; Li et al., 2015), were not changed (Supplemental Dataset 3), suggesting that these play a minor role in vegetative leaf tissue. The SA increase in the absence of UGT76B1 was weaker when UGT74F2 was present (i.e. for the *ugt76b1* and *ugt76b1 ugt74f1* mutants) than when UGT74F2 was absent (i.e. for the *ugt76b1 ugt74f2* and *ugt76b1 ugt74f1 ugt74f2* mutants). Since the upregulation of SA synthesis genes was not affected by the presence of UGT74F2 (Supplemental Table 3A, Supplemental Dataset 3), the increased SGE production likely contributes to the relative drop in SA levels, which remarkably does not affect defense responses (Figs. 3A and 5A). Thus, SGE itself may have a relevant role in defense (Fig. 7C).

Other UGTs were also influenced by mutations of SA glucosyltransferases (Fig. 7A). One of them, UGT75B1, can also form SGE *in vitro* (Lim et al., 2002). However, it did not compensate for the loss of UGT74F2 (Fig. 5A) and overall had a minor impact; in fact, it was downregulated in mutants with increased SA biosynthesis (Supplemental Dataset 3). In contrast, these mutants showed an increased expression of *UGT85A1*, which glucosylates the cytokinin *trans*-zeatin (Hou et al., 2004; Jin et al., 2013; Smehilova et al., 2016). This putative inactivation of cytokinins is consistent with a general growth-defense tradeoff in the plant (Huot et al., 2014) and fits to the downregulation of growth processes most prominently observed in *ugt76b1 ugt74f1* (Fig. 4A). Likewise, *UGT87A2* expression was inversely correlated with UGT76B1 presence. Expression of this gene has been associated with adaptation to abiotic stresses and reduced ROS levels (Li et al., 2017), which matches our ROS measurements (Figs. 6C and 7B). Although ROS signaling plays a major role in defense processes (Waszczak et al., 2018), variation in ROS production across the mutants did not only depend on SA. *UGT87A2* was induced most strongly by *ugt76b1 ugt74f2* (Supplemental Dataset 3), where ROS levels did not differ from the wild type in spite of highly elevated SA. The downregulation of *UGT87A2* by *ugt74f1 ugt74f2* potentially favors ROS induction in spite of missing SA induction.

Mutations of SA glucosyltransferases had profound impacts on the plant phenotype. The SA increase due to loss of UGT76B1 provoked increased resistance to bacterial infection and early senescence of the plants (Figs. 2B-C and 7B; Vlot et al., 2009; Guo et al., 2017). Expression of defense genes across all the mutants was consistent with these phenotypes and the measured SA levels (Figs. 3A and 7C). Remarkably, the loss of almost the whole SA glucosylation capacity did not affect the viability of *ugt76b1 ugt74f1 ugt74f2*. This is due to other mechanisms regulating SA homeostasis by conjugation or degradation that were transcriptionally upregulated upon loss of SA glucosyltransferases, in particular upon the loss of UGT76B1. These transcriptional changes include a slight but significant upregulation of the gene encoding SOT12, which sulfonates SA (Baek et al., 2010), and, more importantly, a substantial enhancement of the SA hydroxylation pathway involving S3H (SA 3-HYDROXYLASE), S5H (SA 5-HYDROXYLASE), and UGT76D1 (Supplemental Table 3B, Supplemental Dataset 3, Fig. 7C), leading to 2,3- and 2,5-dihydroxybenzoic acid and their glucosides (Zhang et al., 2017; Huang et al., 2018). In contrast, the SA methylation pathway remained unaffected (*BSMT1*; Supplemental Table 3B; Chen et al., 2003).

## CONCLUSIONS

This study disentangled individual roles and relationships of the three main *Arabidopsis thaliana* SA glucosyltransferases, UGT76B1, UGT74F1, and UGT74F2, in SA homeostasis and demonstrated their combinatorial effects on development and defense processes. According to our comprehensive mutant comparisons, UGT76B1 is essential for restricting SA levels and is influenced both by UGT74F1 and UGT74F2. UGT74F2 is crucial for producing SGE but does not contribute substantially to SAG production. Steady state increase of SGE and decrease of SA does not decrease transcriptional defense responses, suggesting a defense-supportive role of SGE. Both UGT76B1 and UGT74F1 control defense processes and functionally interact in suppression, but UGT76B1 holds the central and unique role as a defense regulator, tightly coupled with restriction of SA biosynthesis and SA catabolism, whereas UGT74F1 has a subordinate role. Moreover, the two enzymes have different specificities when the plant faces stress. UGT76B1 is the primary SA signaling attenuator during biotic stress, whereas UGT74F1 attenuates SA signaling during abiotic stress. The positive and negative associations between UGTs, SA or SA glucoside levels, and gene expression changes uncovered in this work will direct future studies on spatiotemporal regulation of salicylic acid glucosylation and resistance.

## MATERIAL AND METHODS

### Cultivation and sample collection

Plants were grown in a growth chamber (light/dark cycle 10/14 h at 20/16 °C, 80/65% relative humidity, light at 130 μmol m^-2^ s^-1^) on a peatmoss base (Floragard Multiplication substrate, Germany) and quartz sand substrate mixture (8:1). Three-week-old plants were sprayed with water or BTH (BION™, Ciba-Geigy, Germany) containing 0.01% Silwet L-77 (Lehle Seeds, USA) to support wetting of the leaves. After 1 h, plants were covered with a plastic dome. Leaves were harvested after 48 hours. For one biological replicate, leaves of 25 plants were pooled and immediately frozen in liquid N_2_. The homogenized plant material was split into two parts for metabolite and RNA-seq analysis.

### Infections

Leaves of four-week-old plants were inoculated with *Pst* from their abaxial side using a 1 ml needle-less syringe and 5 x 10^6^ colony-forming units (CFU) ml^-1^ in 10 mM MgCl2. Inoculated plants were covered to maintain high humidity. Leaves were harvested 48 h after inoculation. For one replicate, bacteria from three leaf discs from three plants were extracted in 500 μl 10 mM MgCl2 + 0.01% Silwet L-77 (Lehle Seeds, USA) for 1h and plated for colony counting. Statistical analyses were performed in R 3.6.3 using lm and anova (R Core Team, 2020) and the multcomp package for posthoc testing (Hothorn et al., 2008). Plots were created with gplots (Warnes et al., 2020).

### Staining

GUS histochemical staining performed according to Lagarde et al. (1996). For superoxide analysis, two-week-old plantlets from liquid culture were vacuum-infiltrated with 0.1% (w/v) nitroblue tetrazolium (NBT; Sigma-Aldrich, Germany). Relative saturation was quantified on images of single leaves and compared between groups by nonparametric testing (Supplemental Methods; R Core Team, 2020).

### Metabolic analysis

SA, SAG and SGE levels were extracted from negative mode LC-MS analysis performed on an Ultimate 3000RS (ThermoFisher, Germany) coupled to Impact II with Apollo II ESI source (Bruker Daltonic, Germany; Supplemental Methods).

### RNA-seq analysis

RNA extraction was carried out with the *innuPREP* RNA Kit (Analytik Jena, Germany). All samples had RQN values larger than 8.5 and 260/280 nm absorption between 2.0 and 2.2. Sequencing was performed by BGI Tech Solutions Co., *Ltd*. (Hongkong) using the BGISEQ-500 platform. After alignments against the genome (Berardini et al., 2015) using hisat2-2.1.0 (Li et al., 2009; Kim et al., 2015) and gene expression quantification via stringtie-1.3.4 (Pertea et al., 2015), differential expression analysis was conducted with DESeq2 (Love et al., 2014), followed by GO enrichment analysis and modeling (Supplemental Methods).

### Accession numbers

The RNA-seq data have been deposited in the ArrayExpress database at EMBL-EBI (https://www.ebi.ac.uk/arrayexpress/experiments/E-MTAB-9300).

## Acknowlegements

We thank Wei Zhang for preliminary work on CRISPR/Cas9 editing. We are very grateful to Burkhard Messner for providing the *UGT74F1::GFP-GUS* and *UGT74F2::GFP-GUS* lines.

## Supplemental Material

**Supplemental Data.** Supplemental Figures 1-3, Supplemental Tables 1-3, and Supplemental Methods.

**Supplemental Dataset 1.** GO enrichment analysis for mutant versus wild type comparisons.

**Supplemental Dataset 2.** GO enrichment analysis for functional interactions between UGT enzymes.

**Supplemental Dataset 3.** Significant up- or downregulation of genes for mutants and BTH-treated plants.

